# The Glycosphingolipid GM3 Modulates Conformational Dynamics of the Glucagon Receptor

**DOI:** 10.1101/2020.03.12.988576

**Authors:** T. Bertie Ansell, Wanling Song, Mark S. P. Sansom

**Affiliations:** Department of Biochemistry, University of Oxford, South Parks Road, Oxford, OX1 3QU

## Abstract

The extracellular domain (ECD) of Class B1 G-protein coupled receptors (GPCRs) plays a central role in signal transduction and is uniquely positioned to sense both the extracellular and membrane environments. Whilst recent studies suggest a role for membrane lipids in the modulation of Class A and Class F GPCR signalling properties, little is known about the effect of lipids on Class B1 receptors. In this study, we employed multiscale molecular dynamics (MD) simulations to access the dynamics of the glucagon receptor (GCGR) ECD in the presence of native-like membrane bilayers. Simulations showed that the ECD could move about a hinge region formed by residues Q122-E126 to adopt both closed and open conformations relative to the TMD. ECD movements were modulated by binding of the glycosphingolipid GM3. These large-scale fluctuations in ECD conformation that may affect the ligand binding and receptor activation properties. We also identify a unique PIP_2_ interaction profile near ICL2/TM3 at the G-protein coupling interface, suggesting a mechanism of engaging G-proteins which may have a distinct dependence on PIP_2_ compared to Class A GPCRs. Given the structural conservation of Class B1 GPCRs, the modulatory effects of GM3 and PIP_2_ on GCGR may be conserved across these receptors, offering new insights into potential therapeutic targeting.

**Statement of Significance:** The role of lipids in regulation of Class B GPCRs remains elusive, despite recent structural advances. In this study, multi-scale molecular dynamics simulations are used to evaluate lipid interactions with the glucagon receptor, a Class B1 GPCR. We find that the glycosphingolipid GM3 binds to the glucagon receptor extracellular domain (ECD), modulating the dynamics of the ECD and promoting movement away from the transmembrane domain. We also identify a unique PIP_2_ interaction fingerprint in a region known to be important for bridging G-protein coupling in Class A GPCRs. Thus, this study provides molecular insight into the behaviour of the glucagon receptor in a complex lipid bilayer environment which may aid understanding of glucagon receptor signalling properties.

## Introduction

Class B1 GPCRs are involved in a diverse range of signalling pathways including calcium homeostasis, metabolism and angiogenesis (1). Class B1 receptors are composed of a canonical GPCR seven transmembrane helix bundle (TM1-7), a C-terminal membrane associated helix (H8), and a N-terminal 120-160 residue extracellular domain (ECD). The ECD has a conserved fold (2) and plays a key role in peptide ligand binding, signal transduction and signalling specificity (3). A ‘two-domain’ binding mechanism for peptide ligands has been proposed for Class B1 GPCRs whereby rapid binding of the C-terminus of the peptide to the ECD precedes slower insertion of peptide N-terminus into the transmembrane domain (TMD), leading to conformational rearrangements and receptor activation (4). Differences in the requirement of the ECD for receptor signalling and ligand binding may exist across the Class B1 family. For the Polypeptide-type 1 (PAC1R), Parathyroid hormone (PTH1R) and Corticotrophin-releasing factor 1 (CRF1R) receptors the requirement for the ECD can be bypassed by mass action effects or hormone tethering, consistent with the ‘two-domain’ model and the role of the ECD as an affinity trap. In contrast, for the glucagon receptor (GCGR) and Glucagon-like peptide-1 receptor (GLP1R), the ECD is required for receptor signalling even when the ligand is tethered to the TMD, complicating interpretation of the ‘two-domain’ model (5).

The glucagon receptor (GCGR) is a Class B1 GPCR involved in regulation of glucose homeostasis, amino acid and lipid metabolism (6-8). Consequently the GCGR is a potential candidate for treatment of diseases associated with insulin resistance, such as metabolic syndrome or type 2 diabetes, the prevalence of which increased two-fold over the past 30 years (9). Structures of the full-length GCGR have revealed distinct conformations of the ECD, which differ by rotation around a hinge region linking the ECD to the TMD (10,11). Hydrogen-deuterium exchange experiments alongside MD simulations and suggest the GCGR ECD is mobile and can form transmembrane domain (TMD) contacts in the absence of bound ligand (10,12), further implicating ECD plasticity as a key attribute in GCGR function. Furthermore, a combination of cryo-electron microscopy and MD simulations suggest ECD mobility may be required for binding of peptide ligand to the related glucagon-like peptide-1 receptor (GLP-1R) (13). However, the role of lipids in activation of Class B1 GPCRs is less well understood. Whilst the activation of Class A GPCRs is modulated by membrane lipids (14-16) which may act as allosteric regulators of GPCR activity (15,17,18), the interactions of lipids with Class B GPCRs have not been extensively characterised.

Molecular dynamics (MD) simulations enable exploration of how the physical properties of a membrane (19) and/or direct lipid interactions (20,21) may alter the conformational dynamics of membrane proteins (15,22). For example, a crystal structure of the Class A GPCR β_2_-adrenegic receptor identified cholesterol bound to the intracellular region of TM4 (23), which was validated by observation of cholesterol binding to the same binding site in MD simulations (24). MD simulations have shown how cholesterol binding can modulate the conformation dynamics of the β2 adrenergic receptor (25). MD simulations have also demonstrated that phosphatidylinositol (4,5)-bisphosphate (PIP_2_) binds more favourably to active than to inactive states of the adenosine 2A receptor thus favouring receptor activation (26).

Simulations of GCGR have thus far been limited to bilayers containing just the neutral lipid phosphatidylcholine (PC) (10,12,27,28). Given the role of lipids in GPCR regulation, it is therefore timely to explore the interactions of GCGR with more complex mixtures of lipids, mimicking cellular membranes (29). Furthermore, given the proximity of the ECD to the outer leaflet of the plasma membrane we wished to establish whether an asymmetric and complex lipid environment could influence the dynamics of the ECD relative to the TMD. Using a multiscale MD simulation approach, we combine the enhanced sampling of protein-lipid interactions via coarse-grained (CG) MD simulations with the more detailed representation of interactions in atomistic simulations to probe GCGR dynamics and lipid interactions in in vivo mimetic membrane environments.

## Methods

### Coarse-grained MD simulations

Simulations were performed using GROMACS 5.1.4 (www.gromacs.org). GCGR structures were derived from PDB IDs 5XEZ and 5YQZ (10,11). The T4-lysozyme insert was removed from loop ICL2 and residues between A256 and E260 (5XEZ) or T257 and E260 (5YQZ) were modelled using MODELLER 9.19 (30). The *martinizy.py* script was used to coarse-grain the receptor (31). For the 5YQZ structure the receptor and peptide were coarse-grained separately before consolidation. The ElNeDyn elastic network with a spring force constant of 500 kJ.mol^-1^.nm^-2^ and cut-off of 0.9 nm was applied (32). The transmembrane region of GCGR was embedded in the bilayer using *insane.py* (33) and the receptor centred in a 15 x 15 x 17 nm^3^ box. The system was solvated using MARTINI water (34) and 150 mM NaCl. The system was subject to steepest decent energy minimisation followed by two 100 ns NPT equilibration steps with restraints applied to all protein beads during the first step and just to backbone beads during the second step.

CG simulations were run for 10 μs with a 20 fs integration timestep using the MARTINI 2.2 force-field to describe all components (35). Five or ten repeat simulations of the GCGR structures (5XEZ or 5YQZ) in combination with a bilayer composition as specified in Table 1 were performed, totalling 700 μs of CG simulation data. Temperature was maintained at 323 K using the V-rescale thermostat (36) and a coupling constant τ_t_ = 1.0 ps. Pressure was maintained at 1 bar using the Parrinello-Rahman barostat (37), a coupling constant τ_p_ = 12.0 ps and a compressibility of 3 × 10^−4^ bar-1. The reaction field method was used for Coulomb interactions with a cut-off value of 1.1 nm. VDW interactions were cut-off at 1.1 nm using the potential-shift-Verlet method. The LINCS algorithm (38) was used to constrain bonds to their equilibration values.

**Table 1:**
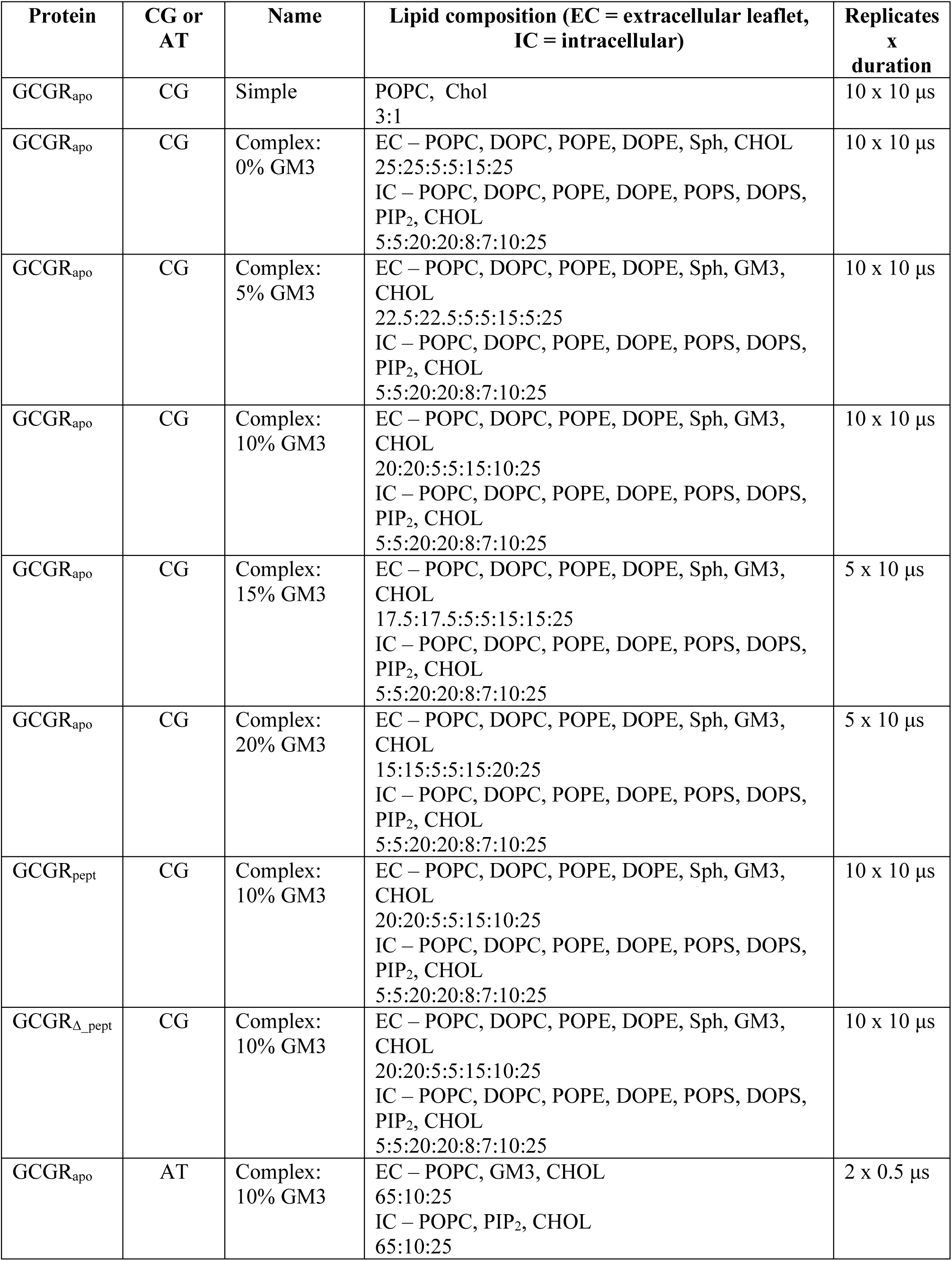
Summary of Simulations.

Protein-lipid interactions were analysed using an in house procedure (PyLipID – https://github.com/wlsong/PyLipID) to calculate the residence time of lipid interactions with GCGR in CG simulations. Briefly, lipid contacts were initiated when the centre of mass of lipid headgroup beads came within 0.55 nm of the protein surface and ended when they exceeded 1.0 nm. Bi-exponential curve fitting of lipid interaction durations as a function of time were used to estimate *k*_*off*_ values for lipid interactions. These *k*_*off*_ values were used to derive lipid residence times which form the basis of the interaction profiles shown in the figures.

### Atomistic molecular dynamics simulations

For atomistic simulations the protein structure was embedded in lipid bilayers which were assembled using the CHARMM-GUI bilayer builder (39). Atomistic bilayers were composed of POPC (65%): GM3 (10%) and cholesterol (25%) in the extracellular leaflet and POPC (65%): PIP_2_ (10%) and cholesterol (25 %) in the intracellular leaflet. The GROMACS 4.6 *g_membed* tool (40) was used to embed GCGR in a bilayer before solvation using TIP3P water (41) and 150 mM NaCl. Steepest decent energy minimisation followed by 5 ns NVT and NPT equilibration steps were performed with restraints applied to the protein.

Two 500 ns atomistic simulations were run for each initial protein conformation, to total 3 μs of atomistic data (Table 1; also see Supporting Material Fig. S1). A 2 fs timestep was used and the CHARMM-36 force-field was used to describe all components (42). Long range electrostatics were modelled using the Particle Mesh Ewald model (43) and a 1.2 nm cut-off was applied to van der Waals interactions. A dispersion correction was not applied. Temperature was maintained at 323 K using the Nosé-Hoover thermostat (44,45) with a coupling constant τ_t_ = 0.5 ps. Pressure was maintained at 1 bar using the Parrinello-Rahman barostat (37), a coupling constant τ_p_ = 2.0 ps and a compressibility of 4.5 × 10^−5^ bar^-1^. All bonds were constrained using the LINCS algorithm (38).

All analysis was carried out using GROMACS 5.1 tools (www.gromacs.org) and locally developed scripts. VMD (46) and pymol (47) were used for visualisation.

## Results & Discussion

### GM3 and PIP_2_ are preferentially localised around GCGR

We wished to explore the effect of lipid bilayer composition on ECD dynamics, given the dynamic behaviour of GCGR observed in previous atomistic simulations (10,12,28) and the proximity of the ECD to the extracellular leaflet of the bilayer. We therefore performed CG MD simulations of the apo state of GCGR (GCGR_apo_, corresponding to PDB id 5XEZ; see Fig. 1A and Methods), of the receptor with a bound glucagon analogue and partial agonist peptide NNC1702 (GCGR_pept_, corresponding to 5YQZ) and of the latter state with the NNC1702 peptide removed (GCGR_Δ_pept_). All three structures were simulated in a ‘complex’ and asymmetric lipid bilayer (PC (40 %): PE (10 %): sphingomyelin (15 %): GM3 (10 %): cholesterol (25 %) in the extracellular leaflet; PC (10 %): PE (40 %): PS (15 %): PIP_2_ (10 %): cholesterol (25 %) in the intracellular leaflet), chosen to mimic the composition of the plasma membrane (Fig. 1B) (48). For each simulation condition (see Table 1), 10 replicates each of 10 μs duration were performed. Previous studies of GPCRs have suggested this is sufficient to adequately sample protein-lipid interactions (14).

**Figure 1:**
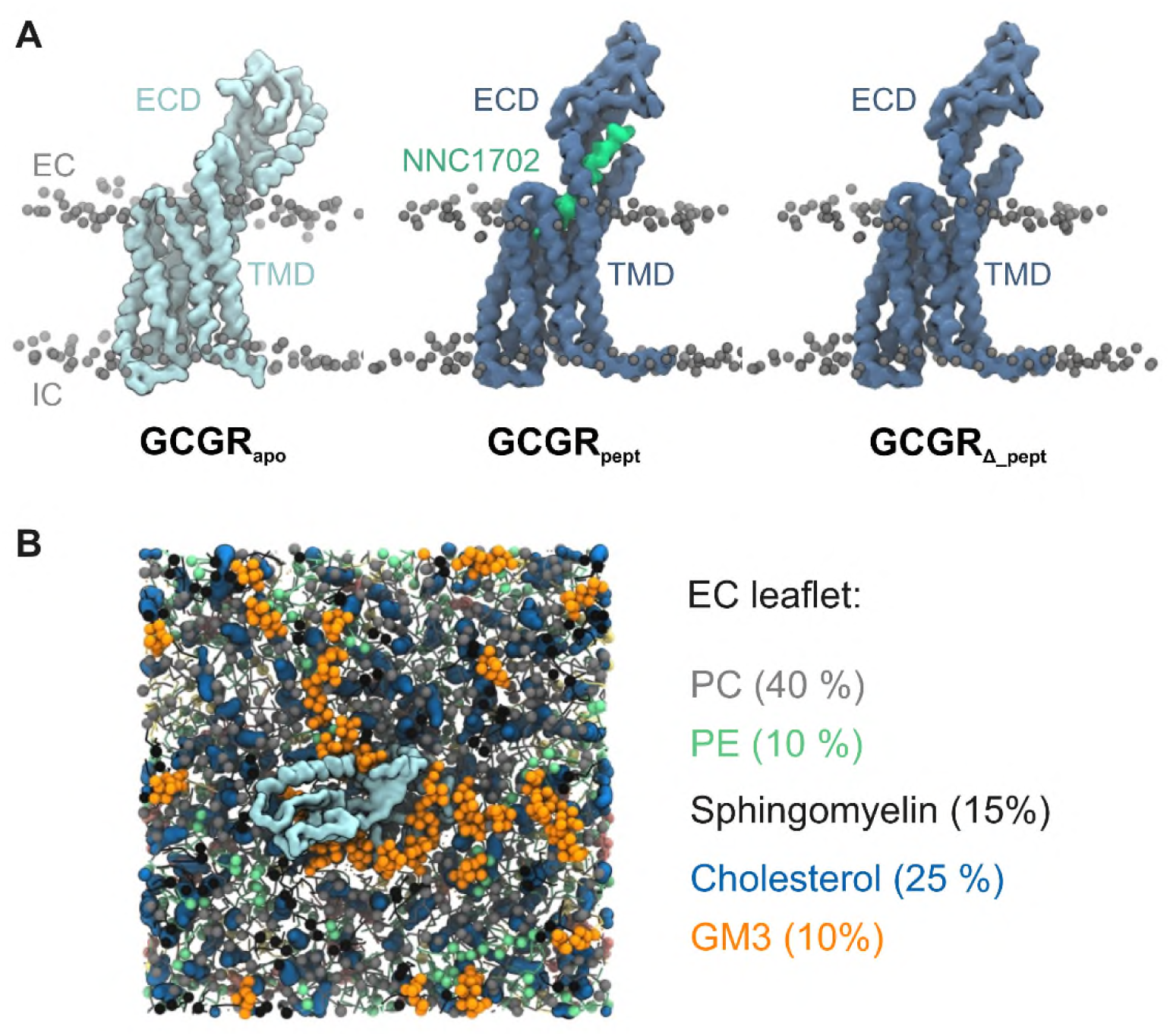
Glucagon receptor (GCGR) structures and lipid bilayer composition. **A)** CG representation of three different GCGR conformations corresponding to GCGR_apo_ (PDB: 5XEZ), GCGR_pept_ (PDB: 5YQZ) and GCGR_Δ_pept_ (PDB: 5YQZ devoid of peptide agonist NNC1702). GCGR backbone beads are shown as quicksurf representations and are coloured light blue (GCGR_apo_) and dark blue (GCGR_pept_/ GCGR_Δ_pept_). Lipid phosphate groups of the extracellular (EC) and intracellular (IC) leaflets are shown as grey spheres and the NNC1702 peptide is coloured lime green. **B)** CG representation of a GCGR_apo_ molecule embedded in a 15 x 15 nm2 ‘complex’ asymmetric bilayer viewed from the extracellular leaflet. Lipids colours are: PC (grey), PE (mint), Sphingomyelin (black), cholesterol (blue), and GM3 (orange).

Comparison of the radial distribution of lipid species surrounding the receptor TMD showed preferential localisation of GM3 and PIP_2_ in the ‘complex’ bilayers (Supporting Material Fig. S2) compared to other lipid species. A locally high radial distribution of GM3 and PIP_2_ has been observed previously for simulations of Class A receptors (26,49) and PIP_2_ binding has been seen during simulations of the Class F GPCR Smoothened (50). Bound PIP_2_ molecules have also been seen in a recent cryo-EM structure of neurotensin receptor 1 (16). However, the current study is the first observation of increased localisation of GM3 and PIP_2_ surrounding a Class B1 GPCR to the best of our knowledge.

### Open and closed conformations of the ECD

Since the ECD of Class B1 GPCRs plays a key role in peptide capture and receptor signal transduction (3) we sought to characterise GCGR ECD conformational behaviour in native-like membranes. This was aided by the two crystal structures of GCGR (5XEZ and 5YQZ) having ECD conformations which differ by an approximately 90° rotation (10,11). The ECD of GCGR_apo_ has a distinct conformation compared to that of GCGR_pept_ which represents the canonical peptide-bound conformation as seen in several Class B1 GPCRs (Supporting Material Fig. S3) (51-58). The stalk in GCGR_apo_ also forms a β-sheet with ECL1. This may be unique to the GCGR apo-state, or may be a consequence of the inhibitory antibody fragment (mAb1) used in crystallisation which binds the ECD and extracellular loop (ECL)-1 (10).

In our CG MD simulations of GCGR in complex membranes (Fig. 2), we observed movement of GCGR_apo_ ECD away from the TMD towards the bilayer (which we will refer to as ECD ‘opening’), around a hinge region formed by residues Q122-E126. This motion permits ECD contact with the bilayer. We also observed movement of the ECD towards the TMD (ECD ‘closure’), consistent with observations in published atomistic simulations in a simple PC bilayer (10).

**Figure 2:**
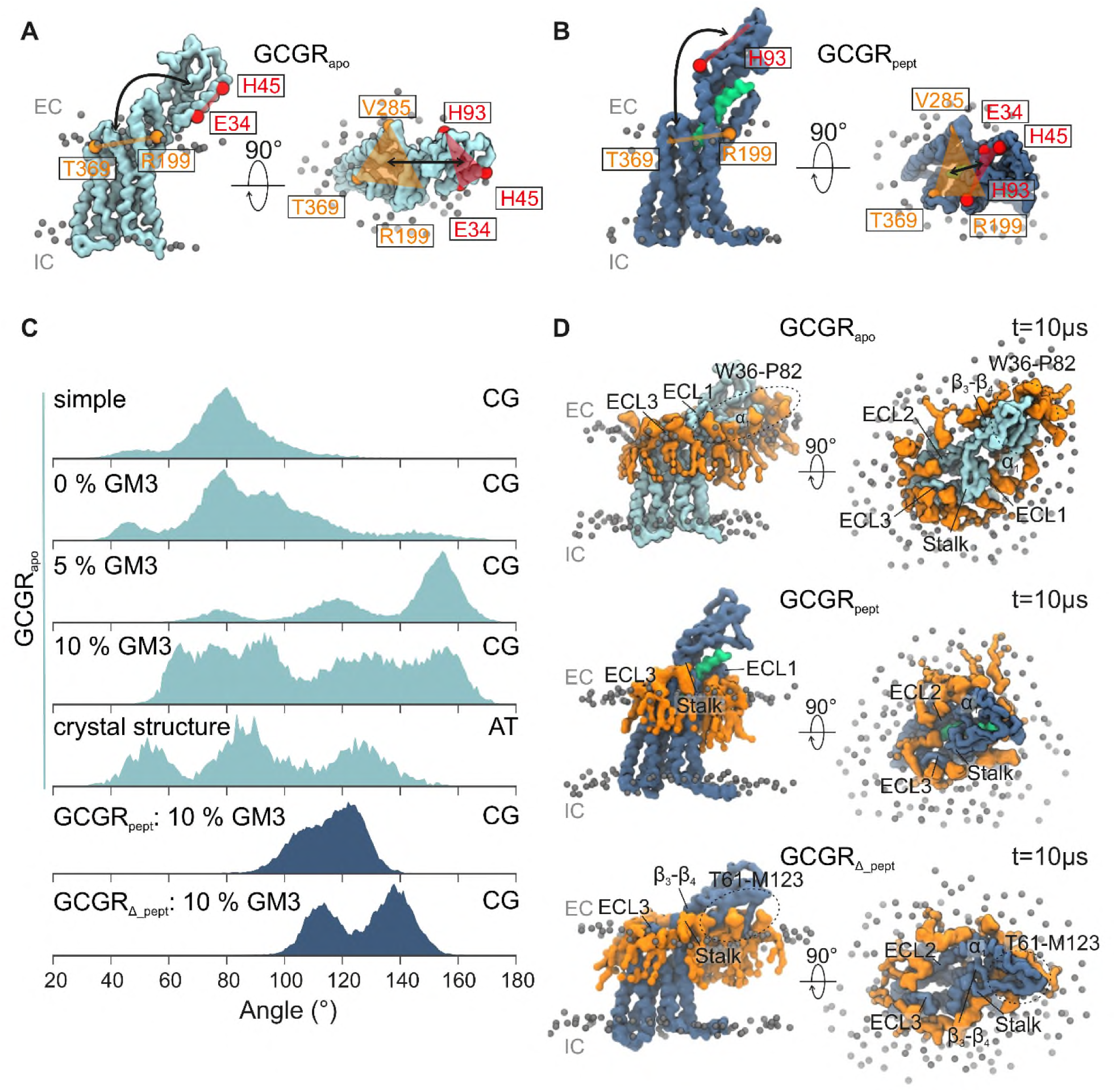
GM3 promotes opening of GCGR ECD towards the bilayer. **A-B)** The angle between two planes (defined by the backbone beads of R199, V285 and T369 on the TMD in orange, and E34, H45 and H93 on the ECD in red) characterises the conformation of the ECD relative to the TMD. These ECD-TMD planes are indicated on CG structures of **A)** GCGR_apo_ (light blue) and **B)** GCGR_pept_ (dark blue). The NNC1702 peptide bound to GCGR_pept_ is coloured lime green. Lipid phosphate groups are shown as grey spheres and the position of the extracellular (EC) and intracellular (IC) leaflets indicated. **C)** ECD-TMD angle distribution calculated across simulations. Simulations from the top down correspond to CG simulations of GCGR_apo_ embedded in a simple POPC:cholesterol bilayer or complex bilayers (as shown in Fig. 1B), containing 0, 5 or 10% GM3, to atomistic simulations of GCGR_apo_ embedded in complex bilayers containing 10% GM3 with the initial protein conformation set to the crystal structure, and to CG simulations of GCGR_pept_ and GCGR_Δ_pept_ in complex bilayers containing 10% GM3. **D)** GM3 (orange) bound to GCGR_apo_, GCGR_pept_ and GCGR_Δ_pept_ at the end of CG simulations in complex bilayers containing 10% GM3. Extracellular loops and regions of the ECD interacting with GM3 are labelled.

Given the increase in bilayer complexity compared to previous simulations, and our observation (above) of GM3 localisation around GCGR, we postulated that ECD opening and/or closing may occur as a result of changes in contacts with the headgroup of the ganglioside (Fig. 2). To investigate the potential effect influence of GM3 on GCGR_apo_ ECD behaviour, we performed CG simulations in both a ‘simple’ bilayer composed POPC:CHOL (3:1) (Fig. 2C & Table 1) and in ‘complex’ bilayers where the abundance of GM3 was modulated between 0 and 10 %, adjusting the amount of PC accordingly (Table 1). We also performed simulations in ‘complex’ bilayers containing an enhanced content of GM3 (15% and 20%, compared to the physiological plasma membrane GM3 concentration of ∼10%; (48)) in order to mimic possible lateral fluctuations in the local GM3 content of cell membranes (59) (Supporting Material Fig. S4A).

To describe motions of the ECD in simulations of the GCGR with different bilayer compositions, we calculated the angle between two planes defined by the residues E34, H45 and H93 on the ECD and R199, V285 and T369 on the TMD (Fig. 2AB). For GCGR_apo_ in either a ‘simple’ bilayer or a ‘complex’ bilayer lacking GM3 the mean (± standard deviation) angle between the ECD and TMD planes was 79° (±16°) and 90° (± 25°) respectively. Inclusion of GM3 in the ‘complex’ bilayer increased the mean angle to 135° (± 27°) and 109° (± 32°) for 5% and 10% GM3 respectively (Fig. 2C). A shift in the distribution of ECD-TMD angles to angles >120° when GM3 is included in the bilayer is consistent with the ability for GM3 to promote a greater range of GCGR_apo_ ECD movement. The increased variability of the ECD-TMD angles when GM3 is included is reflected by a higher standard deviation compared to in the absence of GM3.

Compared to GCGR_apo_, ECD motions were drastically reduced for GCGR_pept_, resulting from peptide contacts bridging the TMD and ECD which restricted domain movement around the hinge region. In the absence of the peptide (simulation GCGR_Δ_pept_) the ECD was observed to move towards the membrane in a manner distinct from that in GCGR_apo_. Visualisation of the trajectories revealed that the ECD closing conformation was maintained by interactions of GM3 with ECD loop W106-A118, and the opening conformation by interactions of GM3 with regions focusing around the α_1_-helix (Fig 2D). We calculated the ECD-TMD angle for GCGR_pept_ and GCGR_Δ_pept_ and, whilst not directly comparable to GCGR_apo_ due to the 90° ECD rotation in the crystal structures, we observed an increase in the mean ECD-TMD angle from 116° (± 10°) for GCGR_pept_ to 128° (± 14°) for GCGR_Δ_pept_ (Fig. 2C). This suggests that for both crystal structures, the ECD conformations are inherently flexible (when devoid of bound peptide) and share a propensity to move towards the membrane. Comparison of the distributions of GM3 around GCGR_apo_, GCGR_pept_ and GCGR_Δ_pept_ at the end of CG simulations in ‘complex’ bilayers containing 10 % GM3 revealed GM3 binding to the ECD of GCGR_apo_ and GCGR_Δ_pept_ but not to GCGR_pept_ (Fig. 2D). This suggests that changes in the conformation of the receptor may be linked to GM3 binding in the absence of bound peptide. Given the structural conservation of the ECD, ganglioside mediated modulation of ECD dynamics might be expected to occur in other Class B1 GPCRs. This in turn could modulate interactions with peptide ligands and/or bias the receptor conformation towards a particular state via sensing of the local bilayer composition.

### ECD movements in atomistic simulations

We have observed ECD movements in multi-microsecond CG simulations, even though an elastic network is present in such simulations (32). To investigate the robustness of these results to the granularity of the simulations, we also performed atomistic simulations (2 × 0.5 μs; Table 1) of GCGR_apo_ starting from the conformation present in the crystal structure. The mean ECD-TMD angle was 92° (± 29 °), i.e. the ECD behaviour in this case showed a mean angle similar to that in CG simulations in ‘simple’ or ‘complex’ bilayers with a low GM3 content. The extent of lipid diffusion during the atomistic simulations allows for just a limited number of GM3 contacts to (re)form with the ECD. Despite this, a peak was observed for ECD-TMD angles > 120 ° (Fig. 2C) and the standard deviation of ECD-TMD angles was high suggesting GM3 has a similar effect on the ECD-TMD angle distribution at atomistic resolution.

In one of the atomistic simulations initiated from the GCGR_apo_ crystal structure we observed closure and subsequent re-opening of the ECD as the simulation progressed (Fig. 3). We analysed the ECD-motions along with GM3 headgroup binding to two regions on the ECD (Site 1 and Site 2) over the course of both atomistic simulations (Fig. 3A-C). In one simulation, GM3 molecules were initially bound to Site 1 on the GCGR_apo_ ECD and the ECD-TMD angle fluctuated around ∼130°. Loss of these GM3 contacts resulted in closure of the ECD towards the TMD (seen as a decrease in angle and increase in RMSD) from 80 ns to 410 ns. GM3 subsequently rebound to Site 1, resulting in re-opening and an increase in the ECD-TMD to 130°. In the second simulation Site 1 was initially occupied by GM3. Again, dissociation of GM3 from Site 1, and subsequent binding of GM3 at Site 2, was accompanied by closure of the ECD. Binding of GM3 at Site 2 locked the ECD in a closed conformation and prevented reopening of the ECD over the course of the simulation. Taken together, these results suggest that interactions of GM3 promote receptor opening, but that this may be modulated by contacts at Site 2 which in turn may favour closure. Furthermore, these observations from the atomistic simulations suggest ECD opening/closure is accessible on the sub-microsecond timescale, and that stable contacts to GM3 at Site 1 are able to maintain an open conformation of the GCGR.

**Figure 3:**
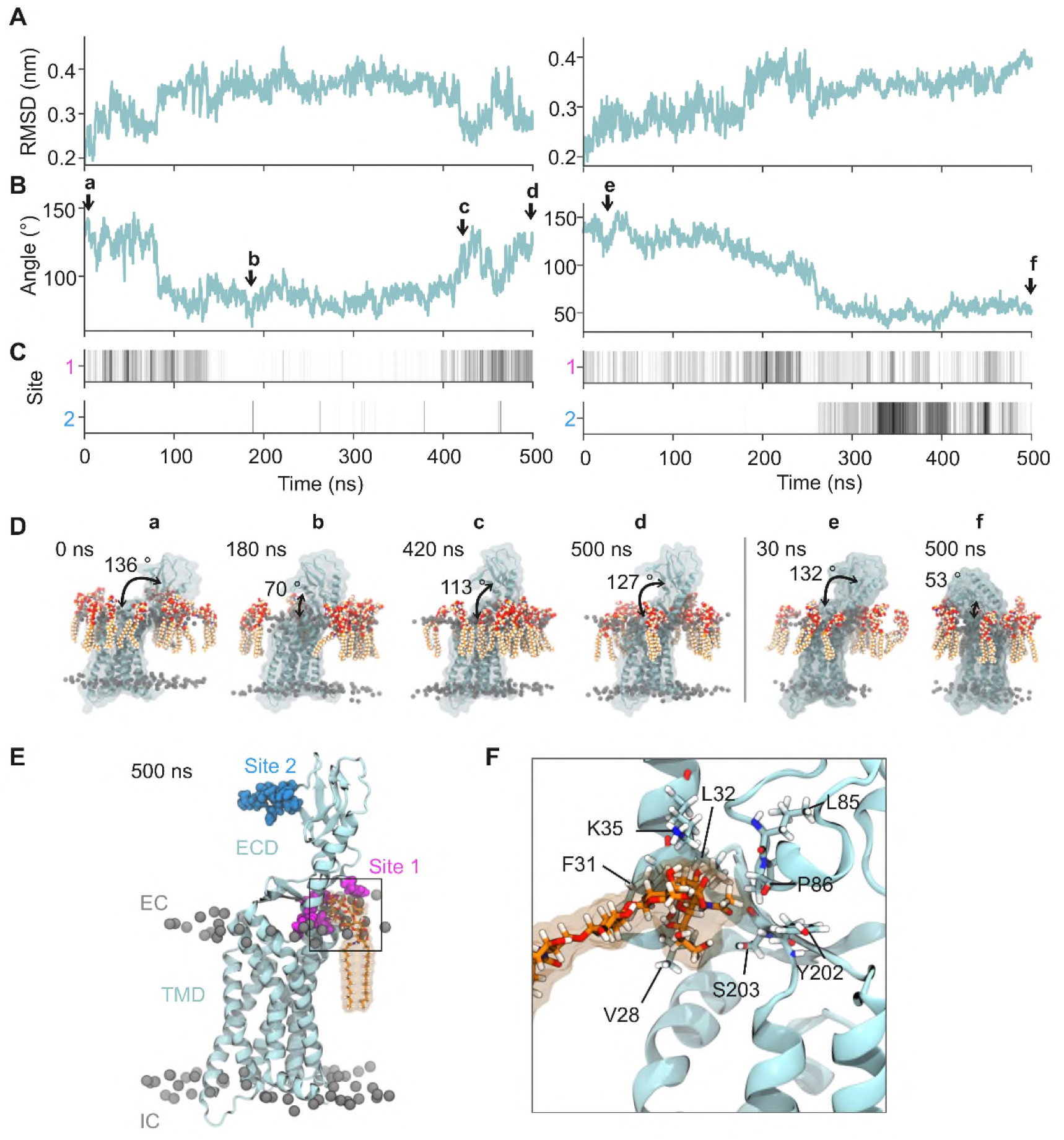
GM3 binding modulates movement of the GCGR_apo_ ECD. **A)** The all atom RMSD of the ECD (residues 27-132) across simulations superimposed on the TMD and (**B**) the ECD-TMD angle changes as a function of time for two 500 ns atomistic simulations of GCGR_apo_ initiated from the crystal structure conformation in the presence of 10% GM3 (see Table 1). In **B** the ECD-TMD angle was defined as the angle between two planes formed by the Cα atoms of R199, V285 and T369 on the TMD and E34, H45 and H93 on the ECD. Arrows indicate snapshots at (**a** to **d**) t = 0, 180, 420 and 500 ns for the first repeat simulation and at (**e** and **f**) t = 30 and 500 ns for the second repeat simulation. **C**) Binding site occupancies for GM3 headgroups within 6 Å of Site 1 and 2 (see parts E/F) over the 2 x 500 ns simulations. Site occupancies were normalised from white (no GM3 headgroups atoms within 6 Å) to black (the maximum number of GM3 headgroup atoms within 6 Å of the site). **D)** Snapshots of GCGR_apo_ from the two atomistic simulations at timepoints corresponding to the arrows embedded in a complex membrane containing 10% GM3 (shown in red/orange). The ECD-TMD angles are marked. **E)** The GCGR at 500 ns showing a GM3 molecule bound to Site 1 on the ECD (residues in pink - see text for further details), and also indicating the location of Site 2 (in blue). **F)** Zoomed in view of GM3 bound at Site 1 indicating the key residues involved in the protein-lipid interactions.

To further compare the conformational dynamics of GCGR in both atomistic and CG simulations in ‘complex’ bilayers containing 10% GM3, we performed principle component analysis (PCA) using trajectories fitted to the TMD (Fig. 4). For GCGR_apo_, the motions of the ECD accounted for by the first principle component were comparable in the CG and atomistic simulations, corresponding to opening and closure of the ECD around the hinge region. The first principle component accounted for 21 to 85 % of the total motion (from the component eigenvalues) in the CG simulations and 23 to 84 % in atomistic simulations. In contrast, for GCGR_Δ_pept_ movement accounted for by the first principle component shows ECD tilting such that the W106-A118 loop approaches the bilayer, accounting for 24 to 63 % of total component eigenvalues. While movement represented by the first principle component of GCGR_pept_ ECD was generally characterised by W106-A118 loop movement towards the bilayer, comparable to GCGR_Δ_pept_, there were small differences in the extent and angle of ECD movement between replicates, suggesting the presence of bound peptide alters the propensity of the ECD to move towards the bilayer. These eigenvalues accounted for 19 to 61 % for GCGR_pept_, slightly lower than those of GCGR_apo_ and GCGR_Δ_pept_.

**Figure 4:**
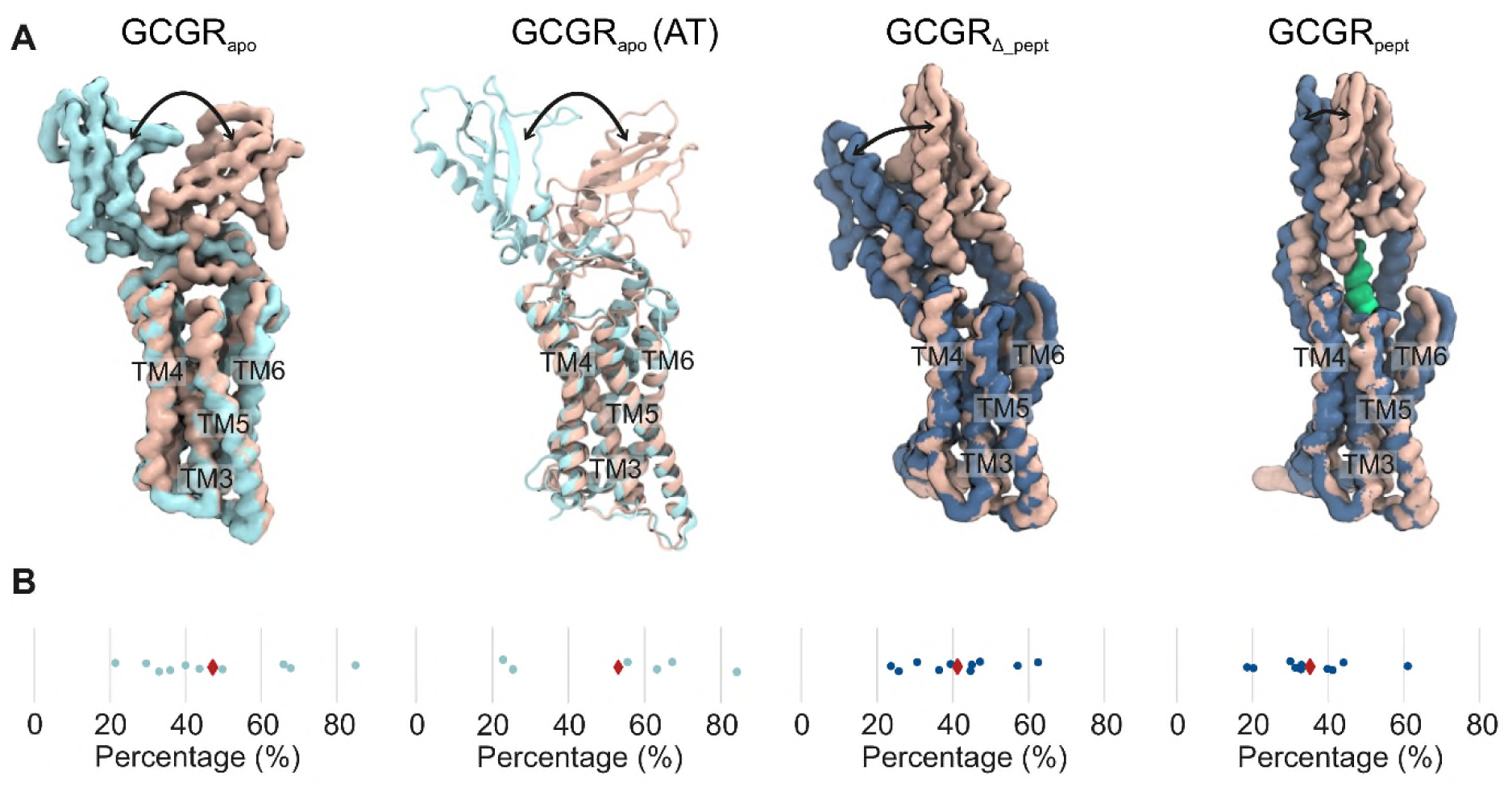
GCGR conformational dynamics. Principal component analysis of GCGR dynamics was performed for CG simulations of GCGR_apo_, GCGR_pept_ and GCGR_Δ_pept_ and for atomistic simulations of GCGR_apo_, all embedded in bilayers containing 10% GM3. **A)** Representative examples of motions corresponding to the first principal component, coloured accordioning to when the ECD is furthest from the bilayer (ochre) or when the ECD opens towards the bilayer (light blue: GCGR_apo_, dark blue: GCGR_pept_/GCGR_Δ_pept_). NNC1702 peptide is coloured lime. Movement of the ECD is indicated by arrows. **B)** The percentages of motion represented by the first eigenvalue for each simulation replicate are shown as blue circles, with the mean percentage for each simulation shown as a red diamond.

Taken together, our results indicate that interactions of GM3 with different regions of the ECD may lead to diverse ECD conformational dynamics. The interactions of GM3 therefore could allosterically modulate the function of GCGR via e.g. altering the rate of ligand recruitment. In our simulations, the hinge region that connects ECD and TMD is flexible, allowing the ECD to adopt different orientations. This flexibility agreed well with the observation of varied ECD conformations among Class B1 GPCRs (Supporting Material Fig. S3), and the importance of ECD dynamics has been stressed in a number of studies (2,13,57,58). Our simulations further reveal that different conformations of ECD have different dynamic behaviour which may have functional relevance, e.g. large-scale movements between closed and open states in of the GCGR may facilitate peptide ligand recruitment to the receptor.

### GM3 interactions with GCGR

Given the observation of close localisation of GM3 around GCGR in the extracellular leaflet, we postulated that GM3 interactions may have a modulatory effect on ECD dynamics. Indeed, a number of recent studies suggest lipids may play a role in the regulation of GPCRs and in coupling to downstream signalling components (14,16,17). We used protein-lipid contact mapping to assess the interaction profiles of GM3 and PIP_2_ with GCGR as a first step towards understanding how these two key lipids might influence GCGR behaviour. GM3 headgroup interactions with the GCGR TMD were conserved across CG simulations in bilayers containing different concentrations of GM3, interacting with ECD loops ECL1-3 and the extracellular regions of TM1-7 (Fig. 5). The GM3 interactions sites seen in atomistic simulations were similar to those in CG, even though less sampling has led to sparser interactions. This good agreement indicates that the observed interactions are consistent between the different simulation granularities.

**Figure 5:**
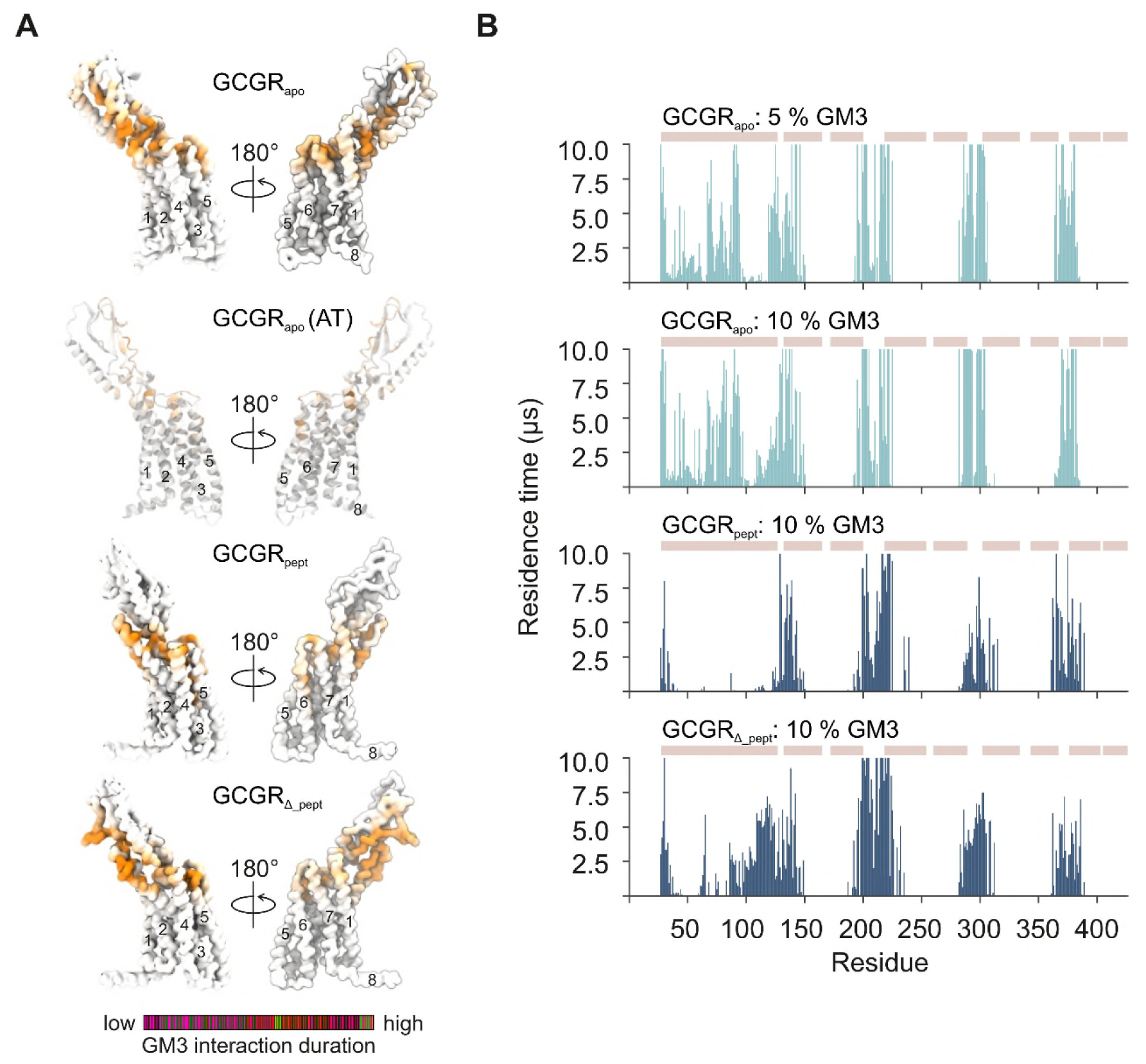
Interactions of GCGR with GM3. **A)** Comparison of GM3 interactions in bilayers containing 10% GM3 mapped onto the structure of GCGR_apo_ from CG and atomistic simulations and GCGR_pept_/GCGR_Δ_pept_ from CG simulations. Contacts are coloured from regions of low (white) to high (orange) mean residence times. **B)** GM3 headgroup interaction profiles with GCGR_apo_, GCGR_pept_ and GCGR_Δ_pept_ in CG simulations in complex bilayers containing 0 - 10 % GM3. GM3 residence times were calculated using a 0.55 nm and 1.0 nm dual cut-off scheme. The position of the ECD, of TM1-7 and of H8 are shown above the contact profile as ochre rectangles.

The GM3 interaction profiles revealed conformational dependence when comparing between the ECDs of GCGR_apo_, GCGR_pept_ and GCGR_Δ_pept_ (Fig.5, Supporting Material Fig. S4B). Thus GCGR_apo_ and GCGR_Δ_pept_ both form interactions of GM3 with the α_1_-helix of the ECD (Q27-K37), with the β_3_-β_4_ loop, and with the ECD-TMD Stalk linker (P86-Q131) due to the proximity of these regions to the bilayer. These interactions overlay with Site 1 discussed above, at which GM3 binding correlates with ECD opening. The interaction fingerprints of GCGR_apo_ at 5% and 10% GM3 are similar, interacting with L38-L85 in addition to ECD regions proximal to the bilayer (Fig. 5, Fig. 2D). In the GCGR_apo_ crystal structure these residues are located 0.5-3.5 nm beyond the terminal GM3 sugar moiety and therefore contacts can only occur when the ECD opens towards the bilayer. For GCGR_pept_ the ECD-GM3 contacts are limited due to restriction of receptor conformation by the bound peptide. GM3 contacts are confined to the α_1_-helix and the Stalk region, within the width of the GM3 glycan layer. When we removed the peptide agonist from our simulations we are able to recover GM3 contacts with D63-D124, including extended interactions with G109-D124 in the GCGR_Δ_pept_ simulations. This suggests that the different ECD conformation do not restrict the ability for the ECD to contact the bilayer but peptide binding does do so.

GM3 binding sites were seen to be more extended than e.g. PIP_2_ binding sites (see below), in part due to the size and flexibility of the ganglioside headgroup. A range of non-polar, polar and positively charged residues interacted with the glycan headgroup. This diversity of GM3 interactions suggests that they may be quite malleable and hence that the observed modulatory effect of GM3 interactions on ECD conformational dynamics could be shared with other Class B GPCRs.

### Differences between PIP_2_ interactions with GCGR and Class A receptors

PIP_2_ has recently emerged as a potential regulator of GPCR state and protein-coupling selectivity (14,16). Analysis of PIP_2_ interactions revealed a conserved interaction fingerprint for all CG simulations of GCGR in a complex membrane (Fig. 6A, Supporting Material Fig. S4C). PIP_2_ molecules bound to sites defined by TM1/ICL1/TM2/TM4, by TM5/ICL3, and by TM6/TM7 and H8, interacting predominantly via their anionic headgroups with cationic (ARG and LYS) residues or via the hydroxyl groups of SER and THR (Fig. 6B). The PIP_2_ contact profile at each interaction site is narrower than for GM3, whilst residence times for PIP_2_ interactions are generally lower than for GM3 (compare Fig. 5B and 6A). For GCGR_apo_, PIP_2_ residence times were longest for the TM1/ICL1/TM2/TM4, TM5/ICL3 and TM6/TM7 sites. GCGR_pept_ / GCGR_Δ_pept_ showed reduced PIP_2_ residence times at TM1/ICL1/TM2/TM4, TM5/ICL3 and TM6/TM7 and enhanced PIP_2_ interaction with H8 compared to GCGR_apo_, suggesting conformation specific differences in PIP_2_ binding, which may be implicated in allosteric regulation of the receptor.

**Figure 6:**
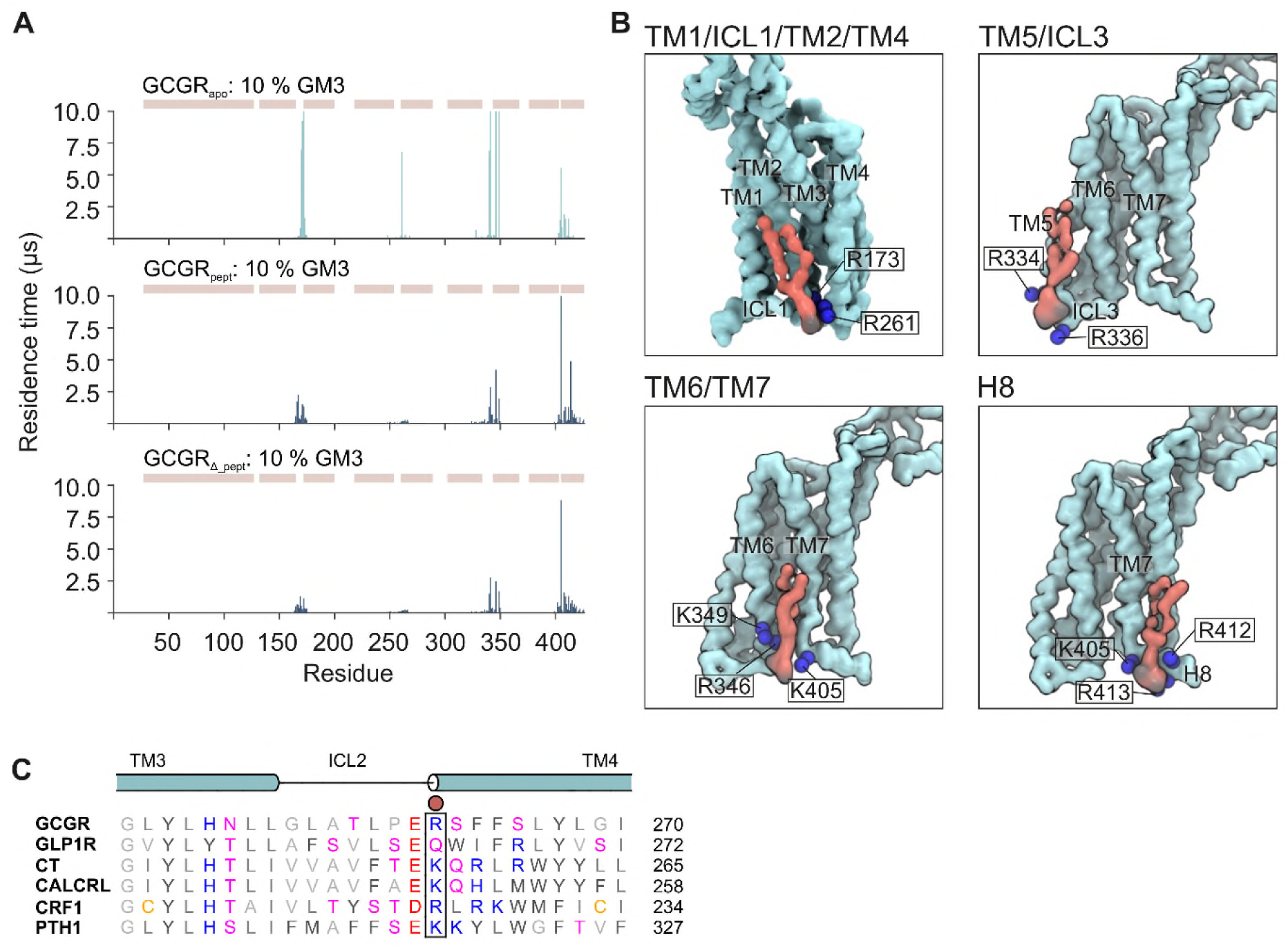
PIP_2_ interactions with GCGR. **A)** PIP_2_ interaction profiles with GCGR_apo_, GCGR_pept_ and GCGR_Δ_pept_ in CG simulations in complex bilayers containing 0 – 10 % GM3. PIP_2_ headgroup residence times were calculated using a 0.55 nm and 1.0 nm dual cut-off scheme. The position of the ECD, TM1-7 and H8 regions are shown above the contact profile as ochre rectangles. **B)** PIP_2_ binding poses identified in CG simulations. PIP_2_ (red) is shown bound to GCGR_apo_ (light blue). PIP_2_ phosphate groups are coloured black and K and R residues are shown as blue spheres. **C)** Structure-based sequence alignment of Class B1 GPCRs showing conservation of the basic R/K residue at the N-terminus of TM4. A red circle shows the position of GCGR_apo_ R261 (see panel **B**) which contributes to binding of PIP_2_ at the TM1/ICL1/TM2/TM4 site. Structure based sequence alignment was performed on GPCRdb.org using the human calcitonin (CT), calcitonin receptor-like (CALCRL), corticotropin-releasing factor 1 (CRF1), glucagon-like peptide-1 (GLP1R), glucagon (GCGR) and parathyroid hormone-1 (PTH1) receptors with manual adjustment based on the position of helices observed in structures.

**Figure 7:**
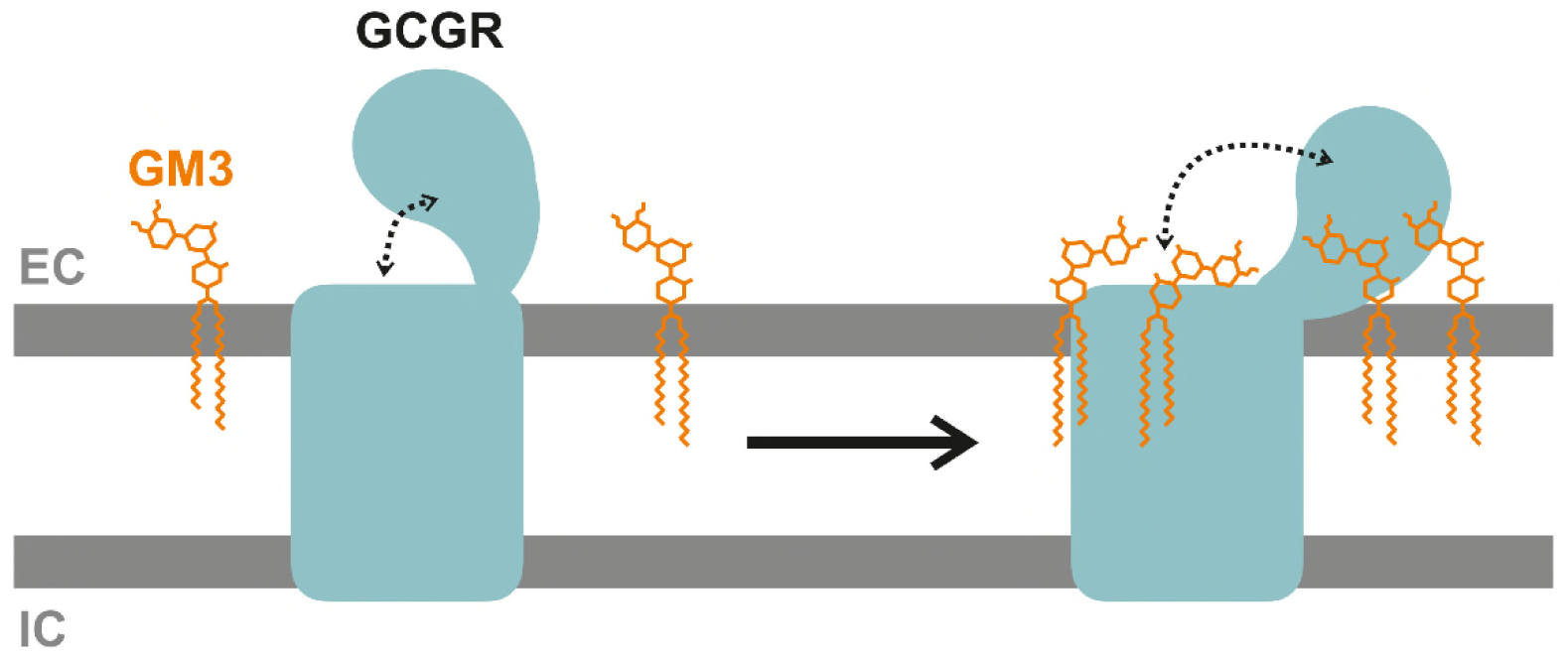
GM3 binding to GCGR promotes ECD opening. Schematic overview of the effect of GM3 (orange) on the behaviour of GCGR (light blue) when devoid of peptide ligands. GM3 binds the receptor TMD and ECD. GM3 binding to the ECD causes conformational modulation of GCGR such that the ECD moves towards the membrane, exposing the peptide ligand binding pocket. The position of extracellular (EC) and intracellular (IC) leaflets are marked.

We compared the interaction profile of GCGR to the prototypical Class A receptor A_2A_ (26). There was agreement between the Class A and Class B1 receptors for PIP_2_ binding to TM1/ICL1/TM2/TM4 and TM6/7. In particular the interaction between the anionic PIP_2_ headgroup and a basic residue at the N-terminus of the TM4 helix (see Fig. 6BC) is conserved across both Class A and Class B1 GPCRs, and is seen in the structure of PIP_2_ bound to the NTS1R/β-arrestin-1 complex (16). However, in contrast to the A_2A_ receptor, PIP_2_ binding was not observed in the vicinity of TM3 or ICL2 of GCGR. Interactions of PIP_2_ with the A_2A_ receptor at TM3/ICL2/TM4 have been suggested to enhance interaction with a mini-Gs-protein, acting as a ‘glue’ between the receptor and the G-protein (26). A lack of PIP_2_ interactions at this site for GCGR may indicate differences in the influence of the anionic headgroups on G-protein coupling that is less dependent on PIP_2_ bridging interactions between the two proteins. Further, a structure based sequence alignment of Class B1 GPCRs showed conservation of positive residues at ICL1 (R/K^12.48^), TM2 (R^2.46^), TM4 (R/K^4.39^), TM5 (K^5.64^, R/K^5.66^), TM6 (R/K^6.37^, R/K^6.40^) and H8 (R/K^8.55^, R/K^8.56^) but not at ICL2 or the intracellular end of TM3 suggesting a lack of PIP_2_ binding at TM3/ICL2 may be a conserved feature across Class B1 GPCRs (Fig. 6C, Supporting Material Fig. S5). This lack of interaction at TM3/ICL2 suggests that the involvement of PIP_2_ in recruitment of signalling partners in Class B1 GPCRs may be different from that in Class A GPCRs.

## Conclusions

MD simulations starting from a number of distinct GCGR conformations have been used to explore the relationship between lipid interactions and the conformational dynamics of the receptor. Two key lipid species, GM3 in the extracellular leaflet and PIP_2_ in the intracellular leaflet, formed contacts with GCGRs. By probing GM3 interactions in different GCGR conformations and in membranes of different GM3 concentrations, we revealed that the binding of GM3 to different parts of GCGR led to generation of different ECD conformations. The multiplicity of ECD conformations could prepare GCGR for the various tasks along its signalling pathways. Given the high degree of structural conservation of ECDs across Class B1 GPCRs (2) and the high level of adaptability to the composition of amino acids in GM3 binding sites, the observed modulatory effect of GM3 interactions on ECD dynamics could be commonplace among Class B1 GPCRs. This could provide a structural explanation for the regulation on GLP1R via localisation to lipid rafts where glycosphingolipids such as GM3 are enriched (60). Indeed, evidence from crosslinking, hydrogen-deuterium exchange, MD and mutagenesis studies suggest that an inactive state of GCGR is favoured by interactions of the ECD with ECL1 or ECL3 (5,10,61). The observation that the binding of GM3 to Site 1 led to opening of GCGR ECD in our simulations suggested that increasing the GM3 concentration in the local environment could shift the receptor towards active states. The varied concentrations of glycosphingolipids in different microdomains of membranes and different compartments of cells could therefore contribute to the multiplication of signalling profiles of Class B1 GPCRs. It is tempting to speculate that changes in lipid metabolism (as a result of dietary intake (62) and/or pharmacological intervention (63)) may lead to changes in lipid rafts. This in turn may affect the relative proportions of the insulin and glucagon receptors localised within raft and non-raft membrane microdomains. GM3 has been observed to promote insulin receptor removal from rafts and decrease insulin signalling (64,65). It is not unreasonable to suggest that GM3 may also play a role in regulation of glucagon signalling and therefore of the overall insulin:glucagon signalling ratio.

In addition to GM3, we identified four PIP_2_ binding sites on GCGR which showed major differences around TM3/ICL2 when compared to PIP_2_ interactions with Class A GPCR A2aR. This could indicate distinct mechanisms of engaging with G-protein and β-arrestin partners whereby Class B1 receptors may have a different dependence on lipid mediatory interactions to bridge the receptor-G-protein interface compared to Class A GPCRs. This may be important for differentiation of receptor signalling and recycling times, potentially contributing to the observation that Class B1 GPCRs have sustained signalling (e.g. (66)) compared to most Class A receptors, postulated to result from enhanced interactions with β-arrestins which may contribute to formation of GPCR G-protein/β-arrestin hybrid complexes (67).

Overall, our simulations provide structural insight into the behaviour of GCGR in complex asymmetric bilayers that mimic the composition of the plasma membrane. We observe modulation of ECD dynamics by the glycosphingolipid GM3, providing an additional layer of complexity to previous observations of GCGR ECD dynamics around the hinge region (10,12). We observe differences in PIP_2_ binding to GCGR compared to Class A receptors which may have functional implications for signalling properties across the Class B1 family. Thus, these data provide a structural basis for further functional investigation of the role of glycosphingolipids and phosphatidylinositols in modulating GCGR signalling, localisation and protein coupling *in vivo*.

## Supporting information

SI Figues S1 to S5

## Acknowledgements

Research in the M.S.P.S. group is supported by Wellcome and BBSRC. W.S. acknowledges support from the Newton International Fellowship. T. B.A acknowledges support from Wellcome. This project made use of time on ARCHER via the HECBioSim, supported by EPSRC.

## Author contributions

M.S.P.S., W.S. and T.B.A conceptualised the project, W.S. and T.B.A. conducted the simulations and analysed the data, and M.S.P.S., W.S. and T.B.A wrote the manuscript.

