## Supplementary material for "The Glycosphingolipid GM3 Modulates Conformational Dynamics of the Glucagon Receptor": SI Figues S1 to S5

**Supporting Material**

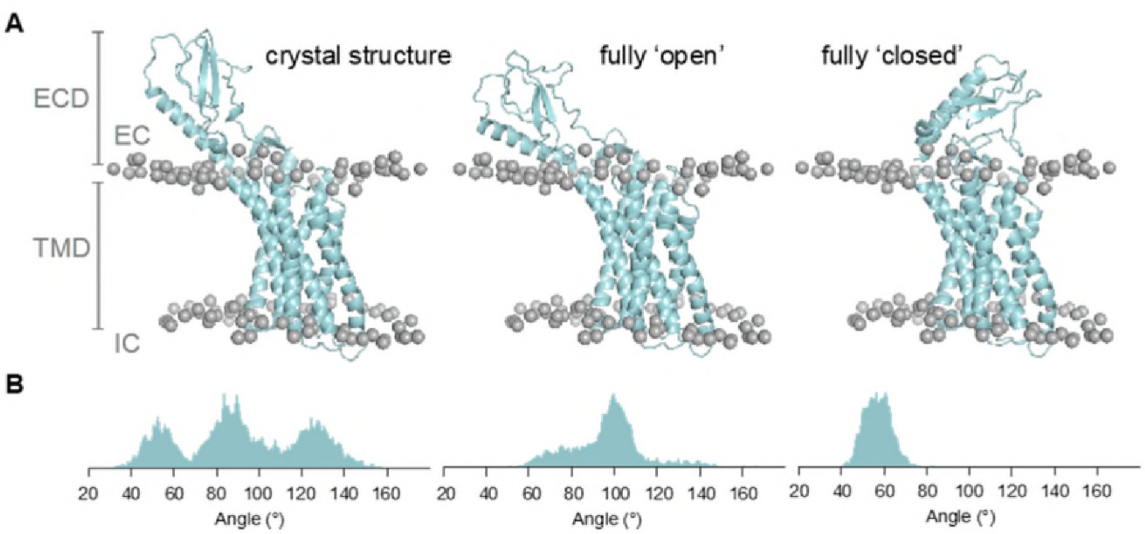


**Supplementary Figure S1: Structures used in atomistic simulations. A)** Comparison of the initial conformations used in atomistic simulations of $GCGR_{apo}$ (light blue). Each conformation was simulated for 2 x 500 ns in a bilayer composed of POPC (65%): GM3 (10%): CHOL (25%) in the extracellular leaflet and POPC (65%): $PIP_2$ (10%): CHOL (25%) in the intracellular leaflet. The fully 'open' and fully 'closed' conformations were backmapped from the end of coarse-grain simulations using the *backward.py* and *initram.sh* scripts. Lipid phosphate groups are shown as grey spheres and the position of the extracellular (EC) and intracellular (IC) membranes are marked. **B)** ECD-TMD angle distribution across each of the simulations setups defined as the angle between two planes formed by the Cα beads of R199, V285 and T369 on the TMD, and E34, H45 and H93 on the ECD.

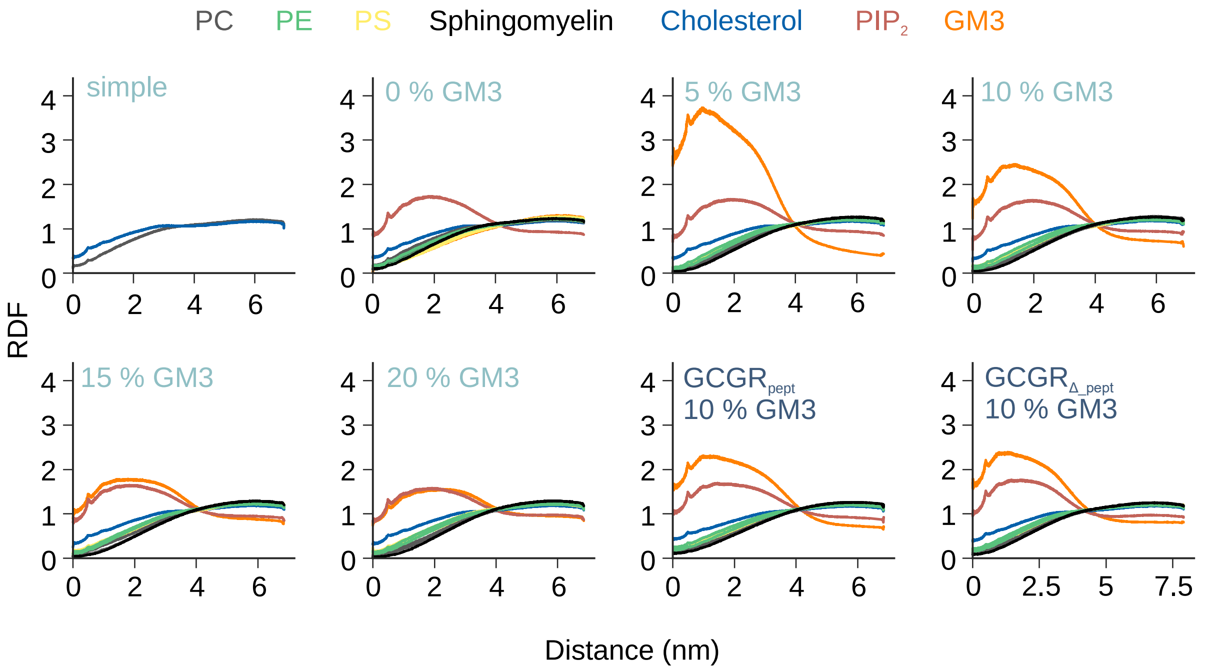


**Supplementary Figure S2: Preferential distribution of GM3 and $PIP_2$ around GCGR.** Radial distribution function (RDF) of lipid species surrounding the GCGR TMD in the xy plane during CG simulations of $GCGR_{apo}$ (light blue), $GCGR_{pept}$ and $GCGR_{\Delta\_pept}$ (dark blue). $GCGR_{apo}$ was embedded in simple or complex bilayers containing (0-20% GM3). $GCGR_{pept}$ and $GCGR_{\Delta\_pept}$ were embedded in a complex bilayers containing 10% GM3. See Table 1 for a detailed description of lipid compositions and simulation times.

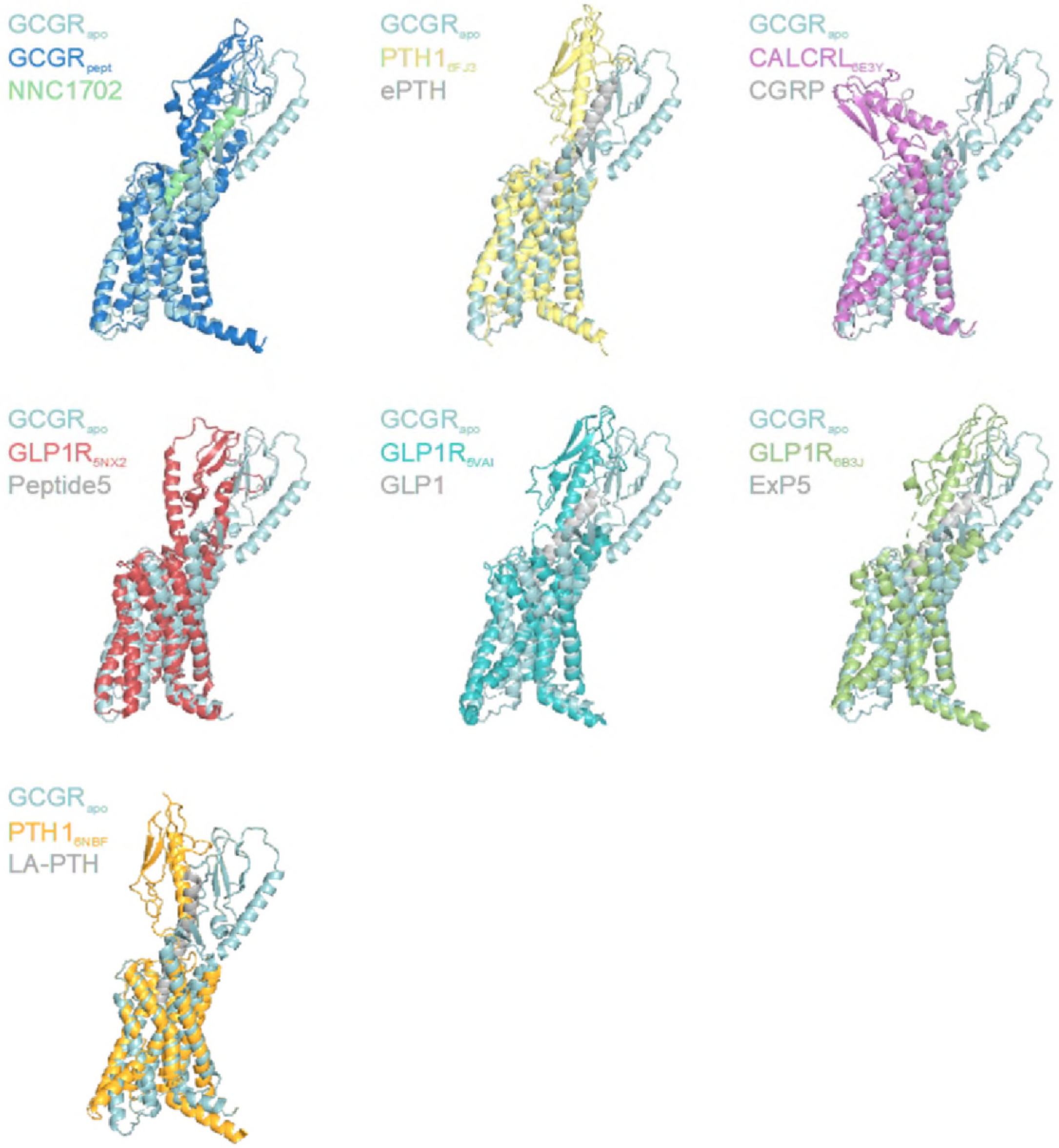


**Supplementary Figure S3: Dynamic conformations of Class B1 GPCR ECDs observed in structures.** Peptide bound multi-domain Class B1 GPCR structures of the glucagon ($GCGR_{pept}$, PDB: 5YQZ), parathyroid hormone 1 (PTH1, PDBs: 6FJ3, 6NBF), calcitonin receptor-like (CALCRL, PDB: 6E3Y) and glucagon-like receptor 1 (GLP1R, PDBs: 5NX2, 5VA, 6B3J) receptors aligned to the TMD of $GCGR_{apo}$ (PDB: 5XEZ) demonstrating the diversity in ECD position observed experimentally. Peptides bound within the TMD pocket are coloured grey with the exception of NNC1702 which is coloured lime green.

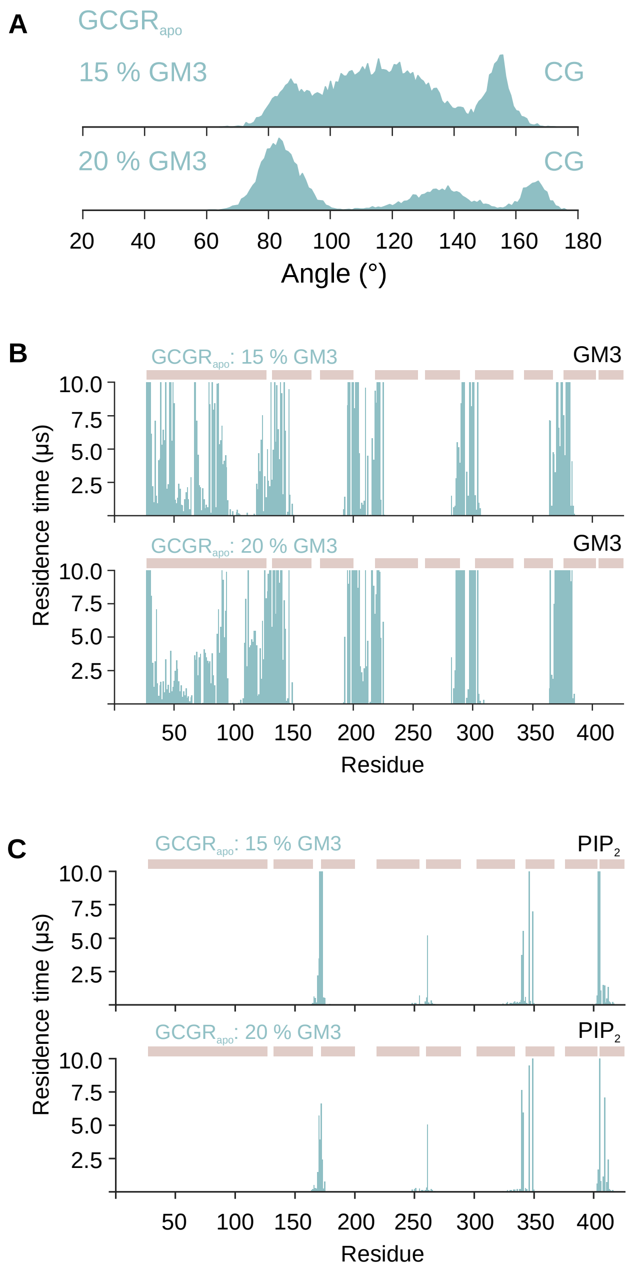


**Supplementary Figure S4: ECD dynamics and lipid contacts in bilayers containing 15% or 20% GM3.** Analysis derived from CG simulations of $GCGR_{apo}$ embedded in complex bilayers containing 15-20% GM3. **A)** ECD-TMD angle distribution across the simulations defined as the angle between two planes formed by the backbone beads of R199, V285 and T369 on the TMD and E34, H45 and H93 on the ECD. **B)** GM3 and **C)** $PIP_2$ headgroup interaction profiles with $GCGR_{apo}$ in CG simulations. Lipid headgroup residence times were calculated using an in house procedure (PyLipID) with a 0.55 nm and 1.0 nm dual cut-off scheme. The position of the ECD, TM1-7 and H8 are shown above the contact profile as ochre rectangles.

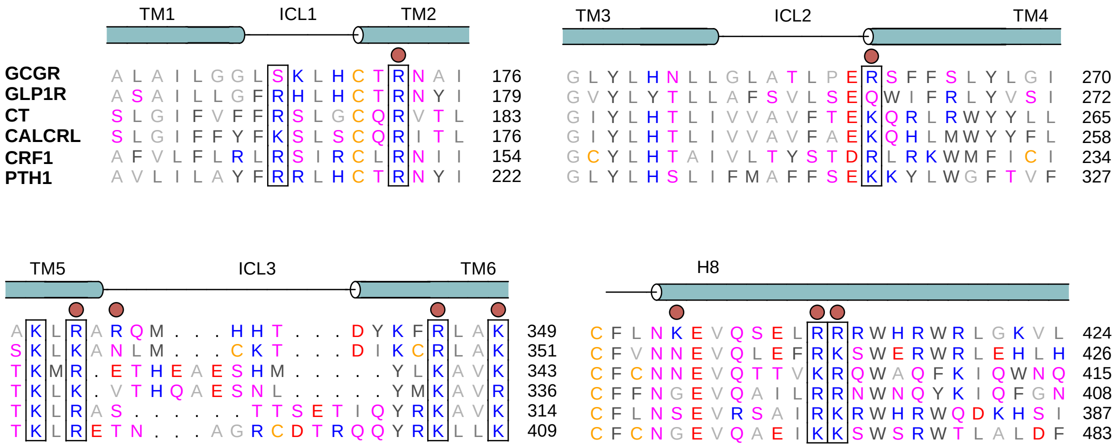


**Supplementary Figure S5: Structure-based sequence alignment of the intracellular regions of Class B1 GPCRs.** Residues are coloured by residue type and the position of helices in the $GCGR_{apo}$ crystal structure indicated. $GCGR_{apo}$ residues which contribute to $PIP_2$ binding are marked by red circles. Black rectangles indicate conserved basic residues. Structure based sequence alignment was performed on GPCRdb.org using the human calcitonin (CT), calcitonin receptor-like (CALCRL), corticotropin-releasing factor 1 (CRF1), glucagon-like peptide-1 (GLP1R), glucagon (GCGR) and parathyroid hormone-1 (PTH1) receptors with manual adjustment based on the position of helices observed in structures.
